# A LOV-domain photoreceptor mediates phototaxis in brown algae

**DOI:** 10.64898/2026.11.07.765074

**Authors:** Carole Duchêne, Susana M. Coelho

## Abstract

Phototaxis enables motile cells to navigate light environments and plays a critical role in survival and reproduction across diverse eukaryotic lineages. This ability has evolved independently through distinct photoreceptor systems. Yet, the molecular mechanisms underlying phototaxis remain poorly understood in many major eukaryotic lineages, including stramenopiles. Here, combining functional genetics, cell biology and evolutionary analyses, we identify Helmchrome (HELM), a LOV-domain protein required for phototaxis in the brown alga *Ectocarpus*. HELM localizes to the posterior flagellum and is responsible for the characteristic flavin-associated fluorescence of brown algal gametes. Loss of HELM abolishes phototactic accumulation and directional orientation without affecting basal motility or fertilization, demonstrating its specific requirement for light-directed movement. Phylogenetic analyses reveal that HELM-like proteins are distributed across diverse stramenopile lineages and that their domain architecture evolved through repeated duplication of an ancestral RGS–LOV module. Our findings identify HELM as a key component of the light-sensing machinery in brown algae, provide a molecular explanation for the long-standing observation of flagellar flavin fluorescence, and reveal a distinct phototaxis system that expands the known diversity of eukaryotic light-sensing mechanisms.

## Introduction

Directional light sensing allows organisms and cells to orient their behaviour relative to a light source and has evolved repeatedly across eukaryotes^1^. In animals, directional photoreception is achieved through diverse arrangements of photoreceptor cells and screening pigments, whereas motile unicellular eukaryotes and unicellular stages of multicellular organisms often rely on compact cellular phototaxis systems. Across microbial eukaryotes, phototaxis is typically mediated by an eyespot, in which pigmented granules interact with a photoreceptor to generate an asymmetric light signal that provides directional information^2–4^. Although this basic organization is broadly conserved, both eyespot architecture and the photoreceptors that mediate light detection differ markedly among eukaryotic lineages. Phototaxis can rely on distinct photoreceptor families, including rhodopsins and flavin-binding proteins, suggesting that different eukaryotic lineages have recruited different molecular solutions for directional light sensing.

One major eukaryotic lineage in which the molecular basis of phototaxis remains poorly understood is the stramenopiles, a diverse group that includes unicellular protists, diatoms and multicellular brown algae. Many stramenopiles produce biflagellate zoospores or gametes with a distinctive light-sensing apparatus, comprising a carotenoid-rich eyespot positioned opposite a swelling at the base of the posterior flagellum, termed the paraflagellar body ^5–7^. This conserved architecture has long implicated the posterior flagellum in directional light sensing, but the molecular mechanism underlying this response remains unresolved.

Brown algae provide a tractable system in which to investigate this mechanism. Their motile gametes exhibit robust phototaxis in response to blue light, with the anterior flagellum providing propulsion and the posterior flagellum contributing to steering^8,9^. Directional illumination induces asymmetric bending of the posterior flagellum, redirecting swimming toward or away from the light source. The close apposition of the paraflagellar body and eyespot, together with strong green autofluorescence of the posterior flagellum under blue-light excitation, pointed to a flavin-based photoreceptor in this compartment ^6^. Consistent with this idea, a unique LOV-domain protein, HELMCHROME (HELM), was identified in association with the posterior flagellum of the brown alga *Colpomenia bulbosa* ^10^. HELM contains two LOV domains predicted to bind flavin chromophores, and these properties made HELM a compelling candidate for the brown algal phototaxis photoreceptor^11^. However, whether HELM actually mediates directional light sensing has not been tested genetically, leaving the molecular identity of the photoreceptor unresolved.

Here, we used CRISPR-Cas genome editing in the model brown alga *Ectocarpus* to determine the function of HELM. We show that HELM localizes to the posterior flagellum and is responsible for the characteristic flavin-associated fluorescence of brown algal gametes. Loss of HELM abolishes directional phototaxis without affecting gamete morphology or fertilization, demonstrating that HELM is the photoreceptor required for light-directed swimming. Our findings identify the molecular basis of phototaxis in brown algae and establish HELM as the founding member of a distinct photoreceptor system that broadens our understanding of how directional light sensing has evolved across eukaryotes.

## Results

### Generation of *helm* knockout mutants in *Ectocarpus*

To investigate the role of HELM in phototaxis, we generated *helm* knockout mutants in *Ectocarpus* using CRISPR-Cas mutagenesis. In the haploid gametophyte, gametes develop within plurilocular gametangia and are released as biflagellate swimming cells (**Figure 1A**). We generated two independent *helm* knockout alleles (**Figure 1B, Figure S1**). All mutations introduced frameshifts leading to premature stop codons, consistent with null alleles (**Figure 1B**).

**Figure 1:**
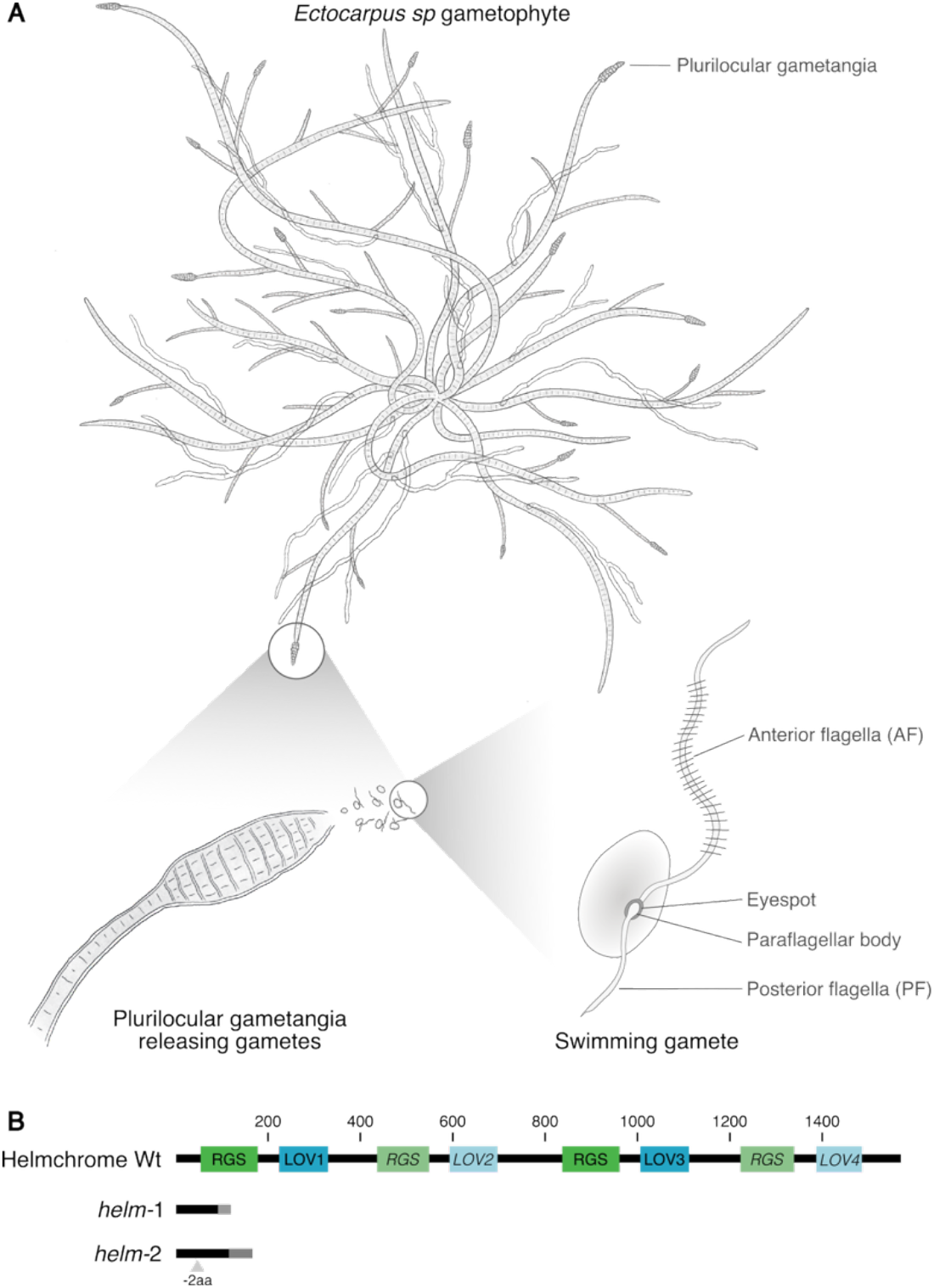
HELM protein architecture and helm mutants. **(A)** Illustration of the biological material used in this study: mature Ectocarpus gametophytes harbor plurilocular gametangia which release swimming gametes. (**B**) Full-length HELM protein and knock-out mutants (helm) generated in this study. LOV: Light-Oxygen-Voltage domain; RGS: regulator of G-protein Signaling domain; Wt, Wild type. Domains in lighter color and italic are thought to be non-functional, see also **Figure S4**; Grey regions in helm-1 and helm-2 indicate non conserved amino acids resulting from frame shifts. See also **Figure S1**.

### HELM accounts for the flavin-associated fluorescence of the paraflagellar apparatus

A characteristic green autofluorescence of the posterior flagellum under blue-light excitation has been observed in brown algal gametes for several decades and attributed to a putative flavin-containing photoreceptor^6^. To determine whether HELM accounts for this fluorescence, we compared wild-type and *helm* gametes.

Swimming wild-type gametes exhibited strong green autofluorescence in the posterior flagellum and paraflagellar body upon blue-light excitation (**Figure 2A**). Strikingly, this fluorescence was absent in helm gametes (Figure 2A), indicating that HELM is required for the characteristic flagellar fluorescence. We further examined freshly settled gametes, in which the flagella are retracted into the cell (**Figure 2B**). In wild-type cells, flagellar retraction produced a diffuse intracellular fluorescence signal with an emission maximum at 520 nm following blue-light excitation, consistent with previous measurements of brown algal flagellar fluorescence^7,12^ (**Figure 2C**). This signal was also absent in *helm* gametes (**Figures 2D, E**). Together, these results demonstrate that the characteristic flavin-associated fluorescence of the posterior flagellum and paraflagellar apparatus depends on HELM.

**Figure 2:**
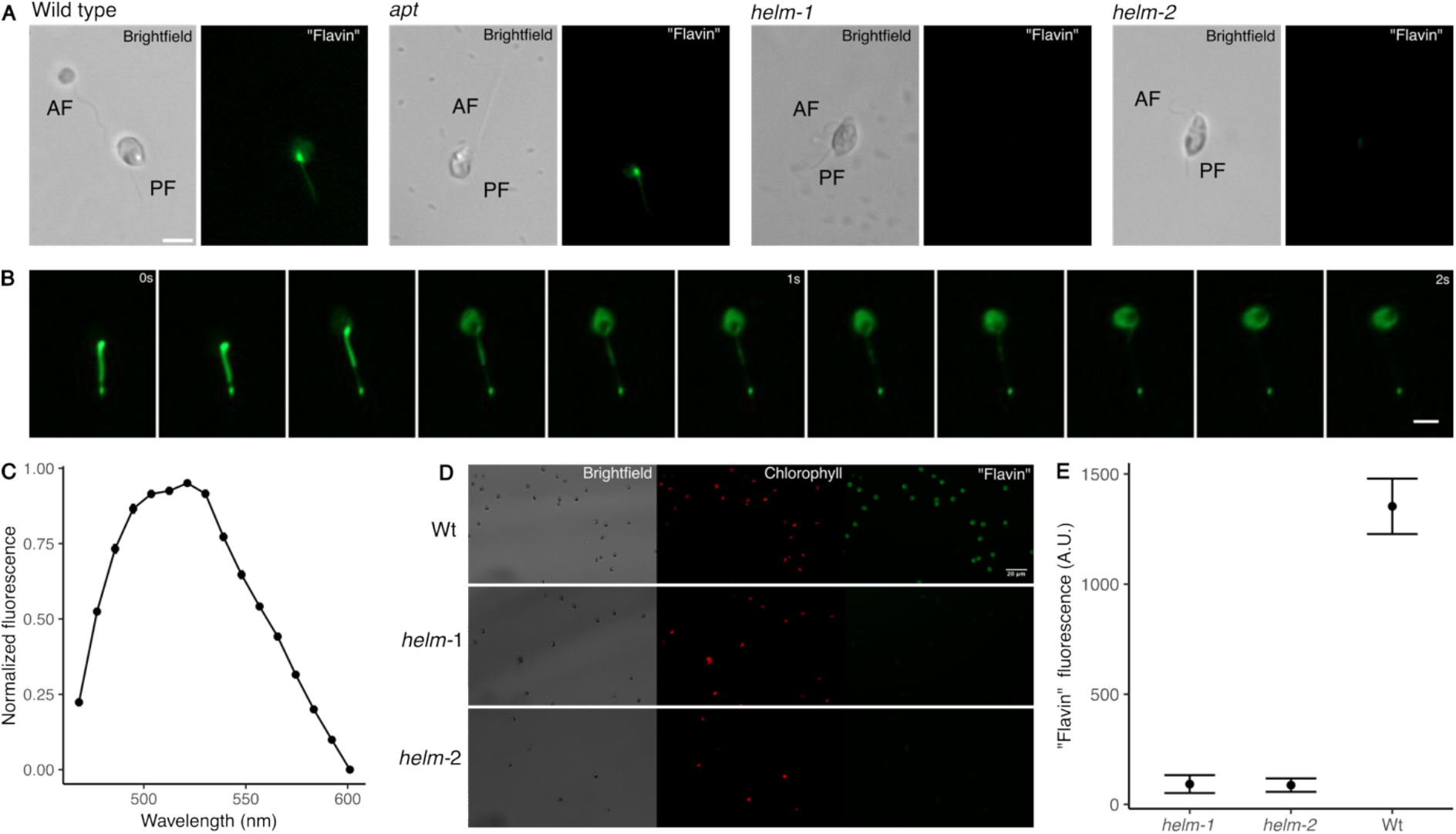
HELM is required for flagellar flavin fluorescence. **(A)** Autofluorescence of swimming wild-type and helm gametes, showing flavin fluorescence in wild-type gametes and its absence in helm gametes. AF, anterior flagellum; PF, posterior flagellum. Scale bar, 5 µm. (**B**) Flagellar retraction during gamete settlement, showing the redistribution of flavin fluorescence to the cell body in wild-type gametes. Images were acquired at 200 ms intervals. Scale bar, 5 µm. (**C**) Flavin fluorescence emission spectra following blue-light excitation in freshly settled wild-type gametes. Mean ± standard error (n = 43 gametes). (**D**) Flavin autofluorescence in settled wild-type and helm gametes. (**E**) Quantification of flavin fluorescence in settled wild-type and helm gametes. Mean ± standard error (n = 20 gametes).

### HELM is required for directional light sensing and phototaxis

To determine whether HELM mediates phototaxis, we first quantified the accumulation of swimming gametes in response to a light source. Wild-type *Ectocarpus* gametes rapidly accumulated in the illuminated region, with a significant increase in cell density detected within minutes of illumination (**Figures 3A–C; Supplementary Video 1**). After 15 min, wild-type gametes showed strong accumulation at the light source, whereas *helm-1* and *helm-2* gametes failed to accumulate (**Figure 3D**), indicating that HELM is required for phototaxis.

**Figure 3:**
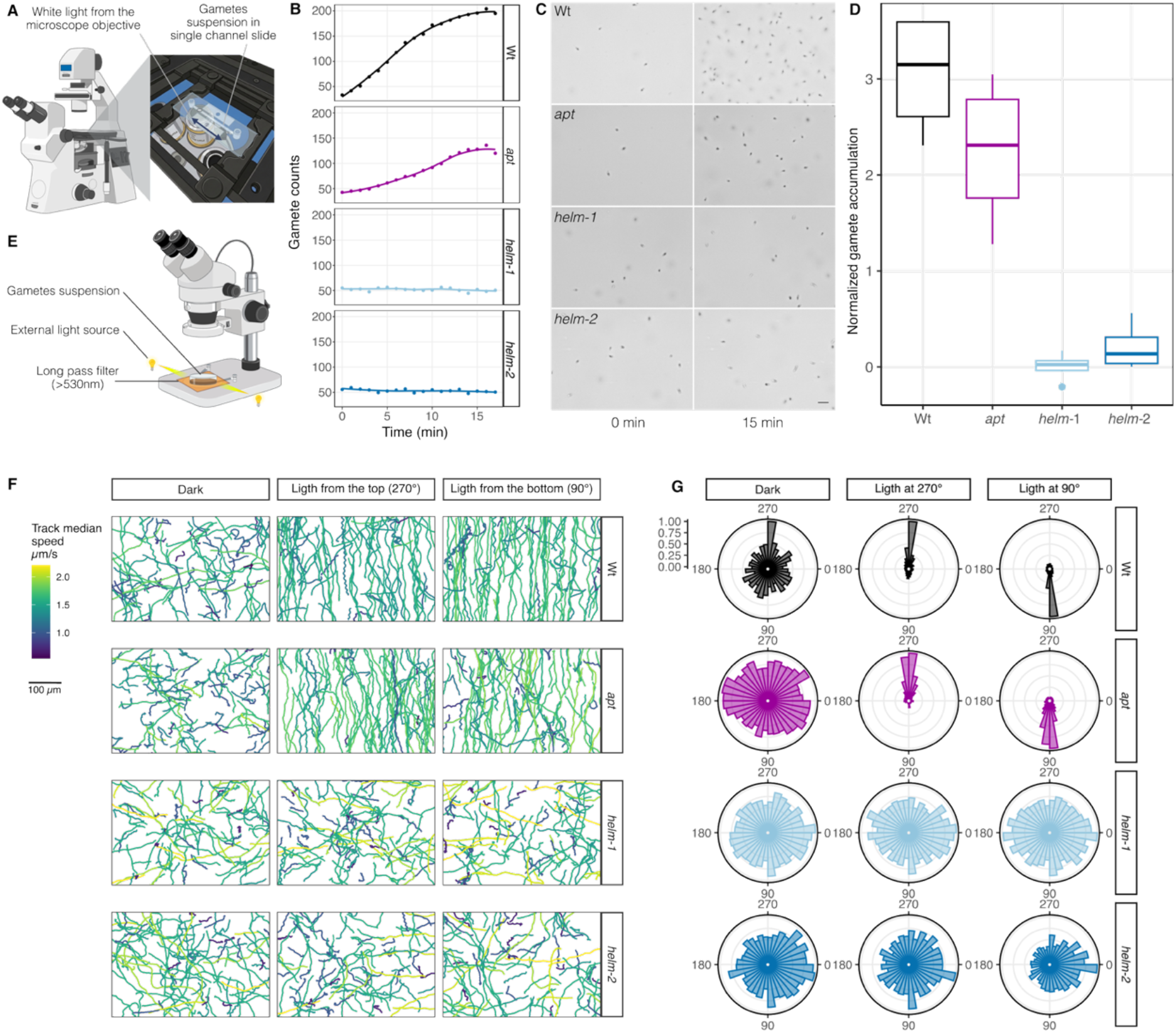
HELM is required for directional light responses. **(A)** Setup for gamete accumulation assay. (**B**) Kinetics of accumulation of gametes towards the light and (C) representative micrographs of accumulation at 0 and 15min (scale bar 10 µm). (**D**) Gamete accumulation after 15min, normalized with the initial gamete concentration at t0. Boxplots are the median of 4 independent biological replicates, each containing 4 gamete accumulation acquisitions per genotype. apt, transformation control (wild-type for the HELM gene, but mutated for the APT gene). (**E**) Setup for gametes swimming direction assay (**F**) 100 randomly selected swimming trajectories in the dark and under directional light; light direction is relative to the horizontal dimension (**G**) Polar histograms of swimming direction under the different light regimes (n>100 swimming trajectories, binned by 10° bin width). A and E were created in BioRender. Duchene, C. (2027) https://BioRender.com/ek4rjr9.

To determine whether the loss of phototaxis resulted from defective directional light sensing rather than impaired motility, we tracked individual gametes under unidirectional illumination (**Figure 3E; Supplementary Videos 2–4**). In darkness, wild-type and helm gametes exhibited comparable swimming behaviour, with no preferred direction, indicating that loss of HELM does not affect basal motility (**Figures 3E and S2**). Upon illumination, however, wild-type gametes rapidly oriented their swimming toward the light source and reversed their direction when the position of the light source was inverted (**Figures 3F and G**). By contrast, *helm* gametes showed no directional bias under illumination and failed to reorient following inversion of the light source. Thus, HELM is required not for swimming itself, but for coupling the direction of incident light to directional movement.

The photobehaviour of wild-type gametes also changed markedly following their release (**Figure S3**). Immediately after release, essentially all photoresponsive gametes displayed positive phototaxis. This relationship progressively reversed: after 3 h, approximately 90% of photoresponsive gametes exhibited negative phototactic behaviour. A similar time-dependent change in photobehaviour has been reported in the brown alga *Mutimo cylindricus*, while mixed populations of positively and negatively phototactic gametes have previously been observed in *Ectocarpus*^8,13^. Together, these results demonstrate that HELM is required for the perception of directional light that underlies phototactic behaviour, rather than for gamete motility itself.

To determine whether loss of HELM causes a general defect in directional responses, we tested the ability of male *helm* gametes to respond to chemical cues from female gametes. Female brown algal gametes release pheromones that attract male gametes by chemotaxis^14^. In the presence of female gametes, wild-type and *helm* male gametes showed comparable directional swimming toward females (**Figure 4A**), indicating that loss of HELM does not impair pheromone-mediated chemotaxis. *helm* male gametes were also able to fertilize female gametes and form zygotes (**Figure 4B**), demonstrating that gamete recognition and fusion remain functional.

**Figure 4.**
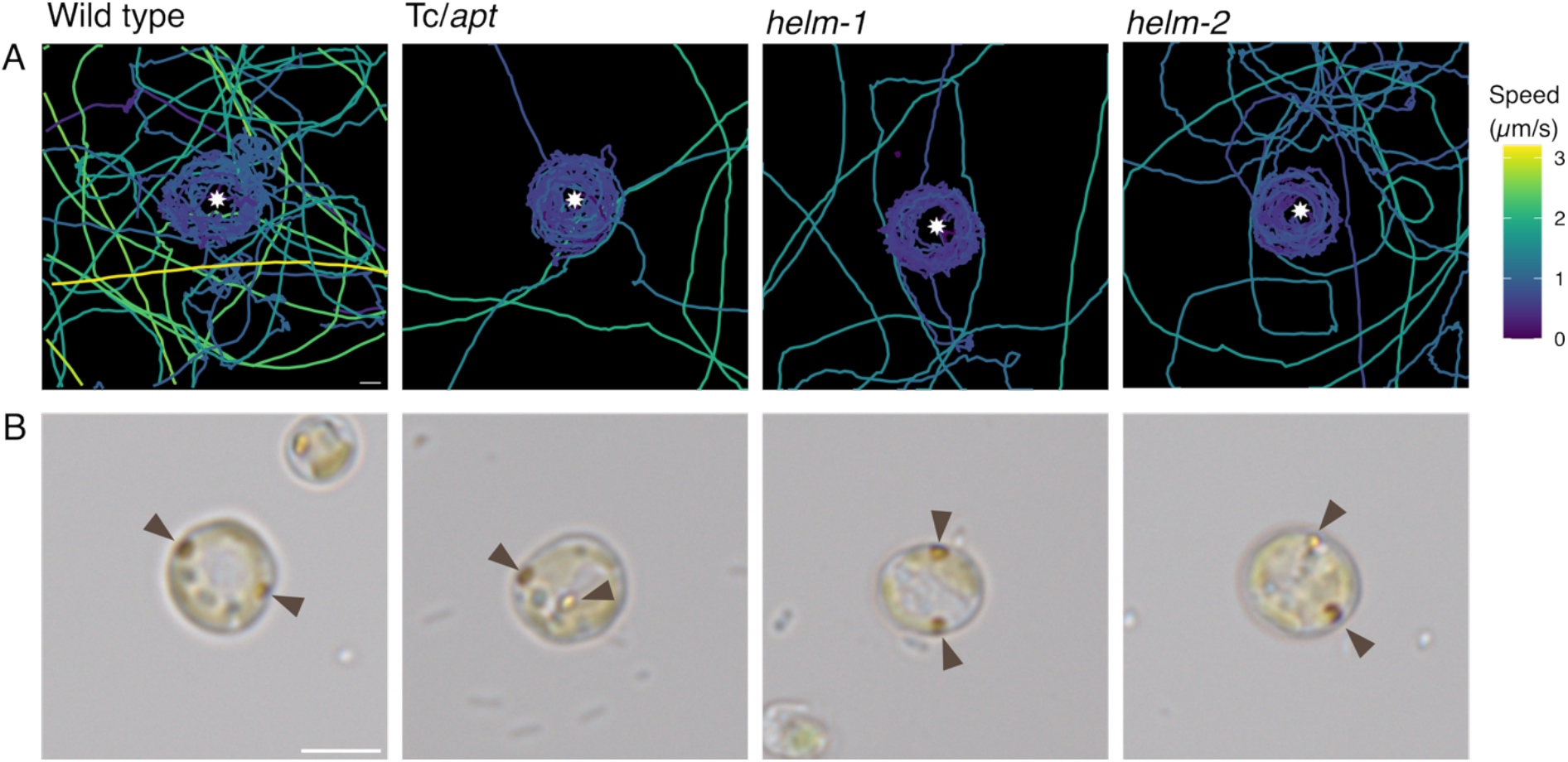
Fertilization is not affected in helm mutants. **(A)** Swimming trajectories of wild type and helm gametes around settled female gametes (asterisk) (**B**) Zygotes observed following the cross of wild type or helm gametes with female gametes. Scale bar, 5µm.

Together, these results show that HELM is specifically required for the response to directional light, rather than for gamete motility, directional swimming or pheromone-mediated chemotaxis.

### HELM evolved from an ancient RGS–LOV protein family

To investigate the distribution and evolutionary origin of HELM, we searched for homologs across Ochrophyta, with an emphasis on brown algae. Previous analyses included only eight brown algal genomes ^15^; we expanded this sampling to 110 brown algal genomes and 42 transcriptomes, together with 184 additional ochrophyte datasets (**Figure 5A-B, Table S1**). Within brown algae, HELM showed a discontinuous phylogenetic distribution. We detected no helm genes in Dictyotales, or in the laminarialean families Laminariaceae, Costariaceae and Alariaceae, a pattern consistent with at least two independent losses of helm (**Figure 5 A**). Across species, the presence of helm broadly correlated with the occurrence of an eyespot in motile gametes or spores, which we used here as a proxy for a potential phototactic system. A notable exception was *Sargassum*: phototaxis, an eyespot and flagellar flavin fluorescence have not, to our knowledge, been reported in Sargassum gametes, yet we identified helm genes or gene fragments in several *Sargassum* genomes. Whether these loci remain functional or represent different stages of HELM pseudogenization remains unclear.

**Figure 5:**
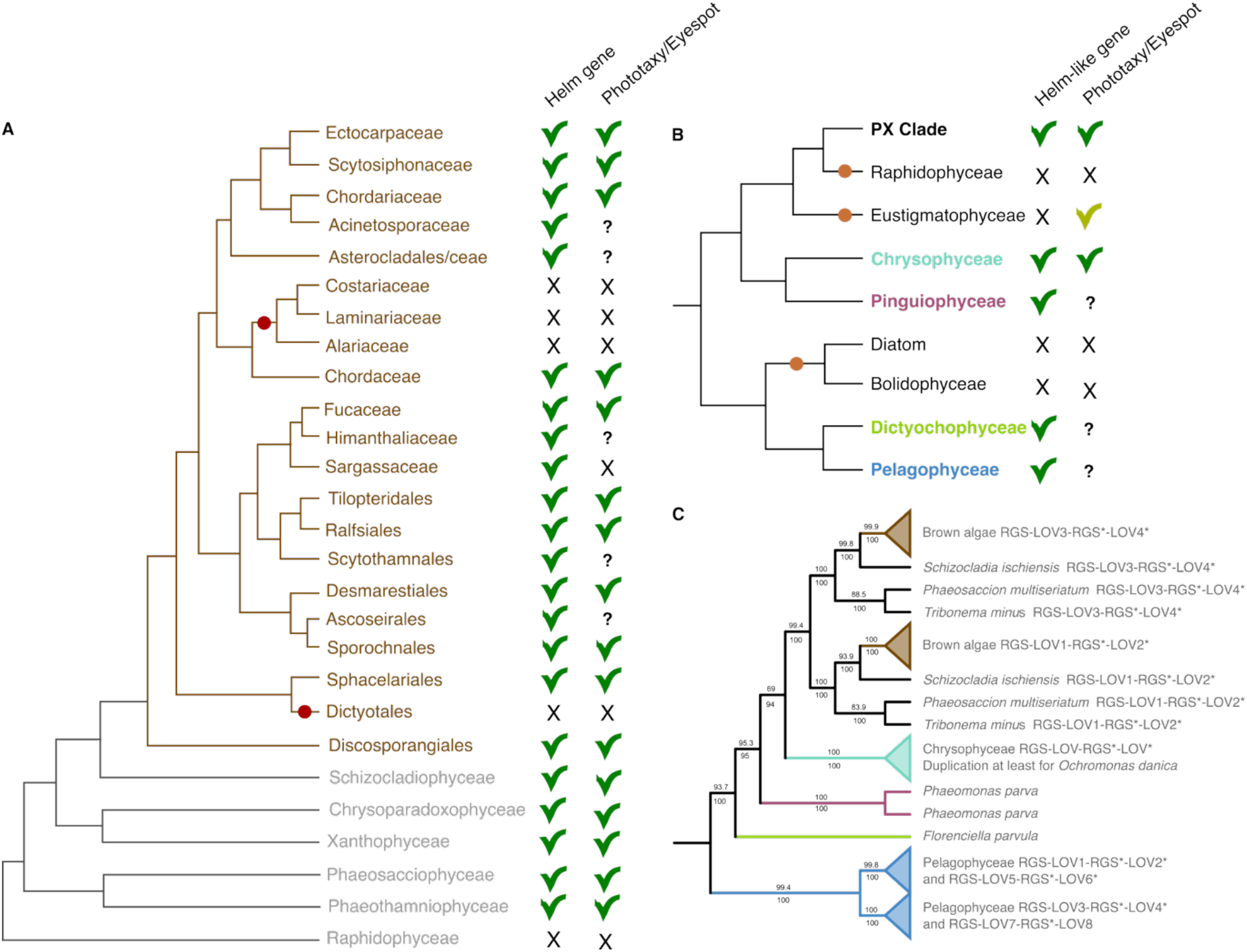
Evolutionary history of the helmchrome gene. A and B, Presence/absence of the helm and helm-like genes in brown algae and sister lineages (A) and across ochrophytes (B). Presence of and eyespot and/or observation of phototaxis was reviewed from the literature (See Table S3); Eustigmatophytes are sometimes reported to possess an eyespot, but structurally different from the eyespots present in other Ochrophytes, hence a different shade of green. C, Phylogenetic tree of the RGS–LOV–RGS*–LOV* repeat. See also **Figures S6B-C** for the phylogenetic tree of the RGS-LOV modules individually. The tree was rooted in Haptophytes HELM -like proteins, which is also supported by the phylogeny of the LOV domain alone (**Figure S6D-E**). numbers above and below each branch are SH-aLRT branch test and ultra-fast bootstraps values, respectively (See Material and Methods).

Importantly, HELM was not restricted to brown algae. We identified bona fide helm homologs in Xanthophyceae, indicating that HELM was already present in the common ancestor of Phaeophyceae and Xanthophyceae (PX Clade, **Figure 5A-B)**. More divergent proteins related to HELM, hereafter HELM-like proteins, were found more broadly across Ochrophyta (Figure 5B), including the Chrysophyceae/Synurophyceae–Pinguiophyceae and Pelagophyceae–Dictyochophyceae clades. Their distribution was again discontinuous, with homologs absent from some sampled genomes, including *Pelagococcus* and *Pelagomonas*.

The occurrence of HELM-like proteins in Chrysophyceae is particularly intriguing. The flagellated chrysophyte *Ochromonas danica* shares several features with brown algal swarmers, including an eyespot facing the paraflagellar body, green flavin-associated fluorescence in the flagellar apparatus, positive HELM-immunostaining and phototactic accumulation in response to blue and green light^6,11,16^. The presence of a HELM-like protein in this lineage therefore raises the possibility that related proteins participate in phototaxis beyond brown algae. By contrast, the role of HELM-like proteins in Pelagophyceae and Dictyochophyceae remains unknown, as phototaxis has, to our knowledge, not been characterized in these groups. Eustigmatophyceae were particularly notable because they possess an eyespot but lack detectable HELM genes. However, their eyespot is structurally different, composed of globular vesicles at the base of their flagella but outside of the chloroplast, indicating that other eyespot-based light sensing mechanisms might exist across Ochrophyta^17,18^.

The patchy distribution of HELM and HELM-like proteins across Ochrophyta could result either from multiple independent origins or from ancestral presence followed by repeated losses. To distinguish between these scenarios, we reconstructed the evolutionary history of their constituent RGS–LOV modules (**Figures 5C** and **S4**). HELM proteins consist of repeated RGS and LOV domains, some of which are strongly diverged. *Ectocarpus* HELM, for example, has the organization RGS–LOV–RGS*–LOV*–RGS–LOV–RGS*–LOV*, where asterisks denote diverged domains predicted to have lost their original function (**Figure S4**). Phylogenetic analysis indicated that the tandem duplication that generated the characteristic HELM architecture occurred in the common ancestor of Phaeophyceae and Xanthophyceae. Independent tandem duplications of related RGS–LOV modules occurred in the Chrysophyceae/Synurophyceae and in the Pelagophyceae lineages (**Figures 5C and S4**), indicating repeated elaboration of the same underlying domain architecture during ochrophyte evolution.

Despite these lineage-specific duplications, the phylogeny of the RGS–LOV modules broadly recapitulated the ochrophyte species tree, with a monophyletic origin, although several relationships had low support and Pelagophyceae and Dictyochophyceae sequences were not recovered as monophyletic (**Figures 5C and S4**). This pattern is most consistent with an ancestral RGS–LOV protein already being present in the common ancestor of Ochrophyta, followed by lineage-specific duplications, diversification and repeated losses. Previous surveys of LOV-domain proteins identified similar RGS–LOV combinations in fungi and haptophytes ^19,20^. In our analyses, fungal RGS–LOV proteins were unrelated to HELM, whereas haptophyte RGS–LOV proteins shared a common origin with the ochrophyte HELM/HELM-like family (**Figure S4 B**). The evolutionary origin of this protein family may therefore predate the diversification of Ochrophyta.

## Discussion

Phototaxis has evolved independently multiple times across eukaryotes, using a common physical principle,i.e. spatially asymmetric light detection, but remarkably diverse cellular architectures and molecular components. In this study, we demonstrate that HELM is required for directional light sensing during phototaxis of brown algal gametes. Loss of HELM abolishes both the characteristic flavin-associated fluorescence of the posterior flagellum and the ability of gametes to orient relative to a directional light source, while leaving swimming and chemotactic responses intact. Together with its localization to the paraflagellar apparatus^11^, these results identify HELM as a central component of the brown algal directional light-sensing system.

HELM represents, to our knowledge, the first LOV-domain protein demonstrated to function in a phototactic system. This contrasts with other characterized eukaryotic phototaxis systems, which employ different classes of photoreceptors, including rhodopsins in green algae such as *Chlamydomonas* and in fungi^2,4^, and a BLUF-domain photoreceptor in *Euglena*^3^. The recruitment of unrelated blue-light receptors to perform equivalent sensory functions illustrates how a common physical requirement, *i*.*e*. determining the direction of incident light, has been solved independently using different molecular components.

An important feature of the brown algal system is that HELM alone cannot encode light direction. Directionality emerges from its spatial organization within the cell: HELM is restricted to the posterior flagellar apparatus, positioned relative to the carotenoid-rich eyespot. Rotation of the swimming cell should therefore generate periodic changes in illumination of the photoreceptor, converting spatial information about the light source into a temporally varying sensory signal. In this sense, directional light perception is an emergent property of the photoreceptor, eyespot and swimming geometry rather than of HELM alone. How light perception is coupled to flagellar movement, potentially through the RGS domains of HELM and other paraflagellar signalling components, remains to be determined.

Our results also separate light perception from the behavioural response it elicits. Newly released gametes are predominantly positively phototactic, whereas older gametes increasingly display photophobic behaviour. As these responses depend on HELM, the same sensory input can therefore generate opposite behavioural outputs depending on gamete state. HELM consequently defines the directional light input rather than the sign of the response, implying that downstream signalling determines whether gametes approach or avoid light. Calcium and cAMP concentrations and presence of female pheromone have been previously identified in *Mutimo cylindrus* as factors involved in the phototactic sign determination^13^. Identifying this switch will be important for understanding how phototactic behaviour is modulated during the reproductive phase of the life cycle.

The evolutionary history of HELM suggests that this photoreceptor did not arise *de novo* in brown algae. HELM homologs are present in Xanthophyceae, placing the characteristic HELM architecture at least in the common ancestor of Phaeophyceae and Xanthophyceae, whereas more divergent HELM-like RGS–LOV proteins occur across several other ochrophyte lineages. Phylogenetic analysis further suggests that an ancestral RGS–LOV protein was already present early in ochrophyte evolution, with the repeated domain architecture characteristic of HELM subsequently arising through lineage-specific duplications. Strikingly, related RGS– LOV modules underwent independent duplications in other ochrophyte clades, suggesting repeated evolutionary elaboration of the same ancestral molecular module. Why this architecture evolved repeatedly is unclear, but the association of a blue-light-sensing LOV domain with an RGS domain provides a potentially direct link between light perception and intracellular signaling. The recurrent duplication of these modules may therefore reflect functional constraints associated with coupling photoreception to rapid changes in flagellar behavior, although this remains to be tested.

The history of HELM also appears to have involved repeated loss. Within brown algae, the absence of *helm* from Dictyotales and several families of Laminariales is broadly associated with the absence of eyespots in their motile reproductive cells, suggesting that loss of the photoreceptor accompanied evolutionary changes in the cellular machinery for directional light sensing. The situation in *Sargassum* may capture a different stage of this process: despite the apparent absence of an eyespot and reported phototactic behaviour, several genomes retain *HELM* genes or gene fragments. Determining whether these loci are expressed and functional, or instead represent ongoing pseudogenization, could reveal how rapidly molecular components are lost following the disappearance of the sensory system in which they function. Conversely, Eustigmatophyceae retain an eyespot but lack detectable HELM homologs, pointing to alternative photoreceptors within Ochrophyta.

The identification of related RGS–LOV proteins in haptophytes potentially push the origin of this protein family deeper still. Haptophytes lack the canonical ochrophyte eyespot, yet flavin-like fluorescence has been reported with an asymmetric distribution between the two flagella in some species^5^. This raises the possibility that the ancestral RGS–LOV protein did not initially function within an eyespot-based phototactic apparatus, but was subsequently recruited into different cellular architectures for directional light sensing. Alternatively, these proteins may have had a more general ancestral role in blue-light signalling and been independently co-opted for phototaxis. Distinguishing between these scenarios will require functional characterization of HELM-like proteins outside the Phaeophyceae–Xanthophyceae lineage.

## STAR Methods

### Resource availability

Mutant lines available upon request to the corresponding authors.

### Experimental model and subject details

All cultured strains are maintained at the Algal Development and Evolution, Max Planck Institute for Biology, Tübingen, Germany. *Ectocarpus sp*.*7* strain Ec32 and mutants generated in this study were cultured in natural seawater (NSW) with half-strength Provasoli solution (0.5 PE) under a 12:12 light:dark cycle (30-40 µmol photon.m^-2^.s^-1^) at 14°C^21^.

## Methods details

### CRISPR-Cas mutagenesis

To generate HELM knock-out mutants, we designed guide RNAs against exons 3 and 4 of the HELM gene (Ec-09_003350) with CRISPOR10^22^ (guide RNA sequences are available in Table S4). For *helm-1* and *helm-2*, transformation and selection were performed as in ^23,24^ using Cas12 enzyme (Alt-R L.b. Cas12a /Cpf1, 15.6 μM, IDT) as follows: Ribonucleoprotein (RNP) were assembled by mixing individual crRNAguide (IDT) at 20 µM final (EcAPT1-Cas12 guide from ^24^ or the guides against the HELM gene, Table S4)) with Cas12 enzyme (1 mg/mL final), in NEB buffer 3.1 (New England Biolabs) (1X final) in IDTE buffer (IDT), incubated for 15min at room temperature, and conserved at 4 °C until use. Gametophytes of the *Ectocarpus sp*.*7* strain Ec32 were grown at 14°C in NSW until maturity (4 weeks) and harvested on a 50µm mesh. Gametophytes were maintained in semi-dry conditions in darkness for 3h at 14 °C, then gametes release was induced by adding ice-cold NSW. Gametes were spin for 10 min at 4600 rpm at 10°C, cell concentration was measured and cells diluted to 1e4 cells/µL. 100 µL of gametes were mixed with RNP mix (12 µL of Helm-targeting RNP, either Cas12-guide1 and 3 or Cas12-guide 2 and 4, plus 6 µL of APT-targeting RNP, 18 µL total) and 120 µL PEG 8000 40% (diluted in NSW) in a petri dish, and left in the dark at room temperature (22°C) for 30min before flooding the plates with 20 mL NSW with half-strength PE. Cells were incubated in the dark at 22°C for 24 h then transferred to 14°C in low light. 2-fluoroadenine was added for selection of mutation in the APT gene 48 h after the transformation, at a concentration of 20 µM. Selection was maintained for 30 days with media refreshment every 10 days before isolation of growing germlings. Putative mutants were screened by PCR (see **Table S4** for primer sequences) and sequenced by Sanger sequencing at Azenta (Genewiz, Germany). An individual that underwent the transformation procedure but did not show mutations in the HELM gene was also isolated (apt control).

### Fluorescence spectroscopy

*Ectocarpus* wild type, *apt, helm-1* and *helm-2* gametophytes were grown at 14°C in NSW+PE until maturity. Gamete release was induced by harvesting gametophytes, quickly drying them with paper, then adding ice-cold NSW. Releasing gametophytes were maintained at 10°C for 30 min, after which gametes were through a 20µm mesh. Swimming gametes were observed in 200µm slide channel (ref) under a Zeiss Axiovert Observer Z1 microscope in brightfield and “Flavin” autofluorescence (excitation LED 469 nm, emission filter 525/50 nm) channels to observe flavin fluorescence. Freshly settled gametes were observed under the same conditions. Fluorescence of 20 individual gametes (and background noise was measured in 10 different positions) were measured with FIJI. Background noise was measured in 10 different positions and subtracted from the fluorescence measure.

For the autofluorescence emission spectra, releasing gametophytes were mounted between a slide and coverslip, and gametes freshly released in the mounted slide were observed with a Zeiss LSM 980 NLO Airyscan microscope. Fluorescence emission was measured every 8nm from 468 to 601 nm under 445 nm laser excitation with a 40X objective with water immersion. Fluorescence was quantified on 43 individual gamete and in 14 positions in the background with the Zeiss Zen software (Zeiss). Background fluorescence was substracted, and fluorescence spectra were normalized to their maximum for each of the 43 individual gametes.

### Phototaxis assays

All phototaxis assay experiments were performed in a dark room at 16°C. For the photoaccumulation assay, gametes were released as explained above and put in a 200µm slide channel (µ-slide 0.2, Luer ibidiTreat, Ibidi). Gamete accumulation was observed on a Zeiss Axiovert Observer Z1 microscope by taking pictures every 5s for 15min at position 1, then moving to position 2 at the other extremity of the slide (2.5 cm distance) for another acquisition, before repeating this sequence (see Figure 3A). The light from the microscope objective was maintained during the whole acquisition. Temperature was maintained at 16°C in the room. Gametes were counted with FIJI ^28^ and differential accumulation was tested by comparing gamete counts during the first and last minute of each acquisition with a Student t.test. While Wt and *apt* gametes always significantly accumulated (p-value <0.05), it was the case only for some technical replicates in *helm* gametes. Gamete accumulation for each acquisition was normalized by doing the difference between gamete counts after 15min compared to t0 divided by the number of gametes at t0. For representation of gamete accumulation with time, gametes counts were smoothed by averaging gametes count over one minute.

For the light direction sensing assay, a long-pass filter (>530nm, Thorlabs) was mounted on an Olympus SZX10 stereomicroscope. Gametes were released as above, and observed in 2mL media in a 35mm petri dish whose sides had been blackened with paint except in 2 positions of about 5mm, opposite to each other, and facing external white LEDs controlled manually. Swimming gametes were observed in the stereomicroscope under dark field (with the long pass filter) under the following sequence: 1min without external light (“Dark”), 1min with external light from one side, 1min with external light from the other side, 1min “Dark”. Movies were acquired at maximal magnification (6.3X) at 70 fps. The resulting movies were analyzed with FIDJ to track gametes using the TrackMate^29^ plugin. Tracks were filtered to remove tracks shorter than 0.5s or with a track displacement smaller than 5 pixels, and slower than 0.7µm/s (to remove floating particles drifting in the media). Swimming direction was assessed by calculating the angle between the start and the end of the track compared to the horizontal dimension.

### Fertilization observations

Female *Ectocarpus sp7* strain Ec25 gametes were released from mature gametophytes as for male gametes and transferred to glass-bottom 35mm dishes (Ibidi). Female gametes were allowed to settle for 1h at 14C in the dark, then the dishes were gently rinsed 3 times with NSW. In parallel, male gametes (Wt, *apt* and *helm*) were released for 30min at 10C. Gametes were observed with a Zeiss Axiovert 5. Male gametes were added at the beginning of the movie and their movement was recorded for 5 min at 70fps. The dishes were then transferred at 14C overnight. On the next day, the dishes were examined for the presence of zygotes, which are distinguished by the presence of two eyespots. For the analysis of male movement around a settled female gamete, an area of ∼100×100 µm centered around a settled female gamete was extracted and male gametes movement were tracked for ∼15s. Tracking was performed using TrackMate^29^.

### HELM and HELM -like proteins in Ochrophyta

An Ochrophyta protein database was built from different genomic and transcriptomic resources (see Table S1). HELM were detected by searching for LOV domains using a custom hidden Markov model (HMM) model built with hmmbuilt from LOV domains from ^19^ .Hmmsearch was run against the Ohcrophyta database and against reference LOV photoreceptors from ^19^, to which we added the Haptophyte RGS-LOV sequences from^20^. All LOV domains were clustered using Sequence Similarity Network (SSN Alignment score 35,^30^), which allowed the separation of helm and helm-like functional LOV domains, nonfunctional ones and LOV domains and other proteins such as Aureochormes in separate clusters. In parallel, a phylogenetic tree containing all LOV domains detected above was built as follows: LOV domain sequences were clustered at 90% identity using cd-hit^31^, aligned using mafft^32^ and trimmed using trimAL^33^ to remove positions conserved in less than 5% of the sequences. The tree was built using IQ-TREE^34^ with automatic model finding, 1000 ultra-fast bootstraps and SH-aLRT branch test (-bb 1000 –alrt 1000). A Helmchrome-specific branch was observed, rooted in Haptophyte sequences. Results of the SSN agreed with the LOV-domain tree, as all HELM and HELM-like sequences identified in the SSN belonged to the same monophyletic clade in the LOV phylogenetic tree. We built trees with RGS-LOV-RGS*-LOV* repeat or the RGS-LOV module using Ochrophyte and Haptophyte sequences. For the different modules, we first performed a repeat analysis on Ectocarpus HELM protein to visualize and define the repetitions with HHrepID ^35,36^. RGS (PF00615) domains were searched for in the 200 amino acids before each HELM and HELM-like LOV and LOV* domain with hmmsearch, aligned with mafft^32^and used to define a HELM-specific RGS domain using hmmbuilt. This new model was used again to select sequences with the RGS-LOV structure, thus defining the RGS-LOV modules. The tree was then built as follows: alignment with mafft^32^, trimming with trimAL^33^ (-gt 0.05) and building with IQ-TREE^34^ (-bb 1000 -alrt 1000) and visualized with Figtree. For the RGS-LOV-RGS*-LOV*, 200 amina acids downstream and 400 amino acids upstream of functional HELM and HELM-like LOV domains were examined for the presence of at least 2 LOV and at least 2 custom RGS modules. Sequences were alignment with mafft^32^ (G-INS-i option), trimming with trimAL33 (-gt 0.1) and building with IQ-TREE34 (-bb 1000 -alrt 1000) and visualized with Figtree and iTOL37. HELM and HELM-like proteins are available in Table S2.

Reference species tree for Phaeophyceae and Ochropytes were drawn from references ^15,38,39^and ^40,41^ respectively. Ability to perform phototaxis and presence of and eyespot was reviewed from the following references ^5–9,11,13,42–47^(see also Table S3).

## Acknowledgements

We thank Taizo Motomura, Chikako Nagasato, Sebastien Colin, Gáspár Jékely, and Christian Feldhaus for fruitful discussions on the phototaxis experimental designs and Daniela (Lulo) Gallego Calderon for the algal drawings and illustrations.

## Author contributions

SMC and CD conceived and designed the experiments. SMC supervised the project and acquired funding. CD performed experiments. CD and SMC curated the data. CD wrote the original draft with support from SMC. SMC reviewed and edited the manuscript. All authors read and approved the final manuscript.

## References

1. Jékely, G. (2009). Evolution of phototaxis. Philos. Trans. R. Soc. B Biol. Sci. 364, 2795–2808. 10.1098/rstb.2009.0072.

2. Kreimer, G., Wakabayashi, K., Hegemann, P., and Dieckmann, C. (2023). The eyespot and behavioral light responses. In The Chlamydomonas Sourcebook (Third Edition), S. K. Dutcher, ed. (Academic Press), pp. 391–419. 10.1016/B978-0-12-822508-0.00004-6.

3. Häder, D.-P., and Iseki, M. (2017). Photomovement in Euglena. In Euglena: Biochemistry, Cell and Molecular Biology Advances in Experimental Medicine and Biology., S. D. Schwartzbach and S. Shigeoka, eds. (Springer International Publishing), pp. 207–235. 10.1007/978-3-319-54910-1_11.

4. Galindo, L.J., Milner, D.S., Gomes, S.L., and Richards, T.A. (2022). A light-sensing system in the common ancestor of the fungi. Curr. Biol. 32, 3146–3153.e3. 10.1016/j.cub.2022.05.034.

5. Kawai, H., and Inouye, I. (1989). Flagellar autofluorescence in forty-four chlorophyll c-containing algae. Phycologia 28, 222–227. 10.2216/i0031-8884-28-2-222.1.

6. Kawai, H. (1992). Green flagellar autofluorescence in brown algal swarmers and their phototactic responses. Bot. Mag. Shokubutsu-Gaku-Zasshi 105, 171–184. 10.1007/BF02489413.

7. Müller, D.G., Maier, I., and Müller, H. (1987). FLAGELLUM AUTOFLUORESCENCE and PHOTOACCUMULATION IN HETEROKONT ALGAE. Photochem. Photobiol. 46, 1003–1008. 10.1111/j.1751-1097.1987.tb04884.x.

8. Kawai, H., Müller, D.G., Fölster, E., and Häder, D.-P. (1990). Phototactic responses in the gametes of the brown alga, Ectocarpus siliculosus. Planta 182, 292–297. 10.1007/BF00197124.

9. Matsunaga, S., Uchida, H., Iseki, M., Watanabe, M., and Murakami, A. (2010). Flagellar Motions in Phototactic Steering in a Brown Algal Swarmer. Photochem. Photobiol. 86, 374–381. 10.1111/j.1751-1097.2009.00676.x.

10. Fu, G., Nagasato, C., Oka, S., Cock, J.M., and Motomura, T. (2014). Proteomics Analysis of Heterogeneous Flagella in Brown Algae (Stramenopiles). Protist 165, 662–675. 10.1016/j.protis.2014.07.007.

11. Fu, G., Nagasato, C., Yamagishi, T., Kawai, H., Okuda, K., Takao, Y., Horiguchi, T., and Motomura, T. (2016). Ubiquitous distribution of helmchrome in phototactic swarmers of the stramenopiles. Protoplasma 253, 929–941. 10.1007/s00709-015-0857-7.

12. Kawai, H., Nakamura, S., Mimuro, M., Furuya, M., and Watanabe, M. (1996). Microspectrofluorometry of the autofluorescent flagellum in phototactic brown algal zoids. Protoplasma 191, 172–177. 10.1007/BF01281815.

13. Kinoshita-Terauchi, N., Shiba, K., Umezawa, T., Matsuda, F., Motomura, T., and Inaba, K. (2019). A brown algal sex pheromone reverses the sign of phototaxis by cAMP/Ca2+-dependent signaling in the male gametes of Mutimo cylindricus (Cutleriaceae). J. Photochem. Photobiol. B 192, 113–123. 10.1016/j.jphotobiol.2019.01.010.

14. Müller, D.G. (1978). Locomotive Responses of Male Gametes to the Species Specific Sex Attractant in Ectocarpus siliculosus (Phaeophyta). Arch. Protistenkd. 120, 371–377. 10.1016/S0003-9365(78)80028-1.

15. Denoeud, F., Godfroy, O., Cruaud, C., Heesch, S., Nehr, Z., Tadrent, N., Couloux, A., Brillet-Guéguen, L., Delage, L., Mckeown, D., et al. (2024). Evolutionary genomics of the emergence of brown algae as key components of coastal ecosystems. Cell 187, 6943–6965.e39. 10.1016/j.cell.2024.10.049.

16. Ishikawa, M., Kataoka, H., and Takahashi, F. (2012). Analysis of Light-Dependent Cell Morphology and an Accumulation Response in Ochromonas danica. Cytologia (Tokyo) 77, 465–473. 10.1508/cytologia.77.465.

17. Hibberd, D.J., and Leedale, G.F. (1972). Observations on the Cytology and Ultrastructure of the New Algal Class, Eustigmatophyceae. Ann. Bot. 36, 49–71. 10.1093/oxfordjournals.aob.a084577.

18. Eliáš, M., Amaral, R., Fawley, K.P., Fawley, M.W., Němcová, Y., Neustupa, J., Přibyl, P., Santos, L.M.A., and Ševčíková, T. (2017). Eustigmatophyceae. In Handbook of the Protists (Springer, Cham), pp. 367–406. 10.1007/978-3-319-28149-0_39.

19. Glantz, S.T., Carpenter, E.J., Melkonian, M., Gardner, K.H., Boyden, E.S., Wong, G.K.S., and Chow, B.Y. (2016). Functional and topological diversity of LOV domain photoreceptors. Proc. Natl. Acad. Sci. U. S. A. 113, E1442–E1451. 10.1073/pnas.1509428113.

20. Coesel, S.N., Durham, B.P., Groussman, R.D., Hu, S.K., Caron, D.A., Morales, R.L., Ribalet, F., and Armbrust, E.V. (2021). Diel transcriptional oscillations of light-sensitive regulatory elements in open-ocean eukaryotic plankton communities. Proc. Natl. Acad. Sci. 118, e2011038118. 10.1073/pnas.2011038118.

21. Coelho, S.M., Scornet, D., Rousvoal, S., Peters, N.T., Dartevelle, L., Peters, A.F., and Cock, J.M. (2012). How to cultivate Ectocarpus. Cold Spring Harb. Protoc. 2012, 258–261. 10.1101/pdb.prot067934.

22. Haeussler, M., Schönig, K., Eckert, H., Eschstruth, A., Mianné, J., Renaud, J.-B., Schneider-Maunoury, S., Shkumatava, A., Teboul, L., Kent, J., et al. (2016). Evaluation of off-target and on-target scoring algorithms and integration into the guide RNA selection tool CRISPOR. Genome Biol. 17, 148. 10.1186/s13059-016-1012-2.

23. Badis, Y., Scornet, D., Harada, M., Caillard, C., Godfroy, O., Raphalen, M., Gachon, C.M.M., Coelho, S.M., Motomura, T., Nagasato, C., et al. (2021). Targeted CRISPR-Cas9-based gene knockouts in the model brown alga Ectocarpus. New Phytol., nph.17525. 10.1111/nph.17525.

24. Martinho, C., Hoshino, M., Raphalen, M., Bukhanets, V., Kerur, A., Bogaert, K.A., Luthringer, R., and Coelho, S.M. (2025). Efficient CRISPR-Cas genome editing in brown algae. Cell Rep. Methods 0. 10.1016/j.crmeth.2025.101273.

25. Nagasato, C., and Motomura, T. (2002). Ultrastructural study on mitosis and cytokinesis in Scytosiphon lomentaria zygotes (Scytosiphonales, Phaeophyceae) by freeze-substitution. Protoplasma 219, 140–149. 10.1007/s007090200015.

26. Fu, G., Nagasato, C., Ito, T., Müller, D.G., and Motomura, T. (2013). Ultrastructural analysis of flagellar development in plurilocular sporangia of Ectocarpus siliculosus (Phaeophyceae). Protoplasma 250, 261–272. 10.1007/s00709-012-0405-7.

27. Reynolds, E.S. (1963). THE USE OF LEAD CITRATE AT HIGH pH AS AN ELECTRON-OPAQUE STAIN IN ELECTRON MICROSCOPY. J. Cell Biol. 17, 208–212. 10.1083/jcb.17.1.208.

28. Schindelin, J., Arganda-Carreras, I., Frise, E., Kaynig, V., Longair, M., Pietzsch, T., Preibisch, S., Rueden, C., Saalfeld, S., Schmid, B., et al. (2012). Fiji: an open-source platform for biological-image analysis. Nat. Methods 9, 676–682. 10.1038/nmeth.2019.

29. Tinevez, J.-Y., Perry, N., Schindelin, J., Hoopes, G.M., Reynolds, G.D., Laplantine, E., Bednarek, S.Y., Shorte, S.L., and Eliceiri, K.W. (2017). TrackMate: An open and extensible platform for single-particle tracking. Methods 115, 80–90. 10.1016/j.ymeth.2016.09.016.

30. Oberg, N., Zallot, R., and Gerlt, J.A. (2023). EFI-EST, EFI-GNT, and EFI-CGFP: Enzyme Function Initiative (EFI) Web Resource for Genomic Enzymology Tools. J. Mol. Biol. 435, 168018. 10.1016/j.jmb.2023.168018.

31. Li, W., and Godzik, A. (2006). Cd-hit: a fast program for clustering and comparing large sets of protein or nucleotide sequences. Bioinformatics 22, 1658–1659. 10.1093/bioinformatics/btl158.

32. Katoh, K., and Standley, D.M. (2013). MAFFT multiple sequence alignment software version 7: Improvements in performance and usability. Mol. Biol. Evol. 30, 772–780. 10.1093/molbev/mst010.

33. Capella-Gutiérrez, S., Silla-Martínez, J.M., and Gabaldón, T. (2009). trimAl: a tool for automated alignment trimming in large-scale phylogenetic analyses. Bioinformatics 25, 1972–1973. 10.1093/bioinformatics/btp348.

34. Wong, T.K.F., Ly-Trong, N., Ren, H., Baños, H., Roger, A.J., Susko, E., Bielow, C., Maio, N.D., Goldman, N., Hahn, M.W., et al. (2025). IQ-TREE 3: Phylogenomic Inference Software using Complex Evolutionary Models.

35. Gabler, F., Nam, S.-Z., Till, S., Mirdita, M., Steinegger, M., Söding, J., Lupas, A.N., and Alva, V. (2020). Protein Sequence Analysis Using the MPI Bioinformatics Toolkit. Curr. Protoc. Bioinforma. 72, e108. 10.1002/cpbi.108.

36. Biegert, A., and Söding, J. (2008). De novo identification of highly diverged protein repeats by probabilistic consistency. Bioinformatics 24, 807–814. 10.1093/bioinformatics/btn039.

37. Letunic, I., and Bork, P. (2024). Interactive Tree of Life (iTOL) v6: recent updates to the phylogenetic tree display and annotation tool. Nucleic Acids Res. 52, W78–W82. 10.1093/nar/gkae268.

38. Heesch, S., Serrano-Serrano, M., Barrera-Redondo, J., Luthringer, R., Peters, A.F., Destombe, C., Cock, J.M., Valero, M., Roze, D., Salamin, N., et al. (2021). Evolution of life cycles and reproductive traits: Insights from the brown algae. J. Evol. Biol. 34, 992–1009. 10.1111/jeb.13880.

39. Akita, S., Vieira, C., Hanyuda, T., Rousseau, F., Cruaud, C., Couloux, A., Heesch, S., Cock, J.M., and Kawai, H. (2022). Providing a phylogenetic framework for trait-based analyses in brown algae: Phylogenomic tree inferred from 32 nuclear protein-coding sequences. Mol. Phylogenet. Evol. 168, 107408. 10.1016/j.ympev.2022.107408.

40. Terpis, K.X., Salomaki, E.D., Barcytė, D., Pánek, T., Verbruggen, H., Kolisko, M., Bailey, J.C., Eliáš, M., and Lane, C.E. (2025). Multiple plastid losses within photosynthetic stramenopiles revealed by comprehensive phylogenomics. Curr. Biol. 0. 10.1016/j.cub.2024.11.065.

41. Cho, A., Lax, G., and Keeling, P.J. (2024). Phylogenomic analyses of ochrophytes (stramenopiles) with an emphasis on neglected lineages. Mol. Phylogenet. Evol. 198, 108120. 10.1016/j.ympev.2024.108120.

42. Kawai, H. (1988). A Flavin-Like Autofluorescent Substance in the Posterior Flagellum of Golden and Brown Algae1. J. Phycol. 24, 114–117. 10.1111/j.1529-8817.1988.tb04464.x.

43. Coleman, A.W. (1988). The Autofluorescent Flagellum: A New Phylogenetic Enigma. J. Phycol. 24, 118–120. 10.1111/j.1529-8817.1988.tb04465.x.

44. Kawai, H., Kubota, M., Kondo, T., and Watanabe, M. (1991). Action spectra for phototaxis in zoospores of the brown algaPseudochorda gracilis. Protoplasma 161, 17–22. 10.1007/BF01328893.

45. Ichinomiya, M., dos Santos, A.L., Gourvil, P., Yoshikawa, S., Kamiya, M., Ohki, K., Audic, S., de Vargas, C., Noël, M.-H., Vaulot, D., et al. (2016). Diversity and oceanic distribution of the Parmales (Bolidophyceae), a picoplanktonic group closely related to diatoms. ISME J. 10, 2419–2434. 10.1038/ismej.2016.38.

46. Wetherbee, R., Jackson, C.J., Repetti, S.I., Clementson, L.A., Costa, J.F., van de Meene, A., Crawford, S., and Verbruggen, H. (2019). The golden paradox – a new heterokont lineage with chloroplasts surrounded by two membranes. J. Phycol. 55, 257–278. 10.1111/jpy.12822.

47. Henry, E.C. (1987). Primitive reproductive characters and a photoperiodic response in Saccorhiza dermatodea (Laminariales, Phaeophyceae). Br. Phycol. J. 22, 23–31. 10.1080/00071618700650031.

