## Supplementary material for "A LOV-domain photoreceptor mediates phototaxis in brown algae": Figure S: SuppFigures_020126.docx

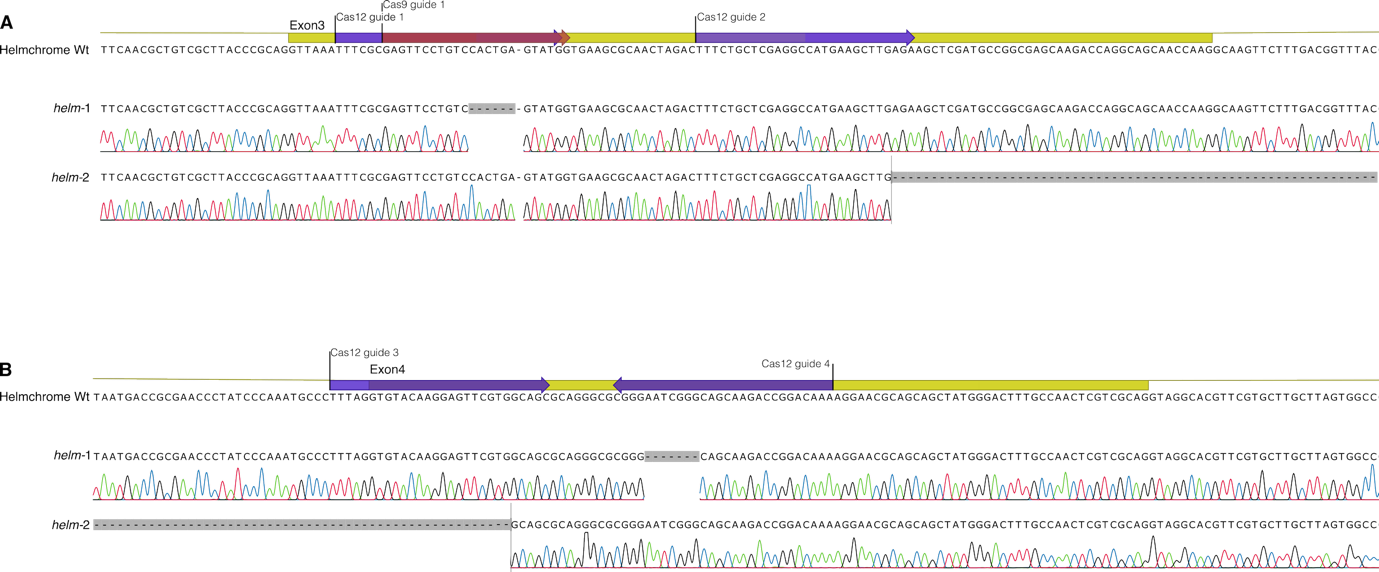

### **Figure S1**: Crispr-Cas induced mutations of the HELM gene in helm strains in Exon3 (A) and 4 (B)

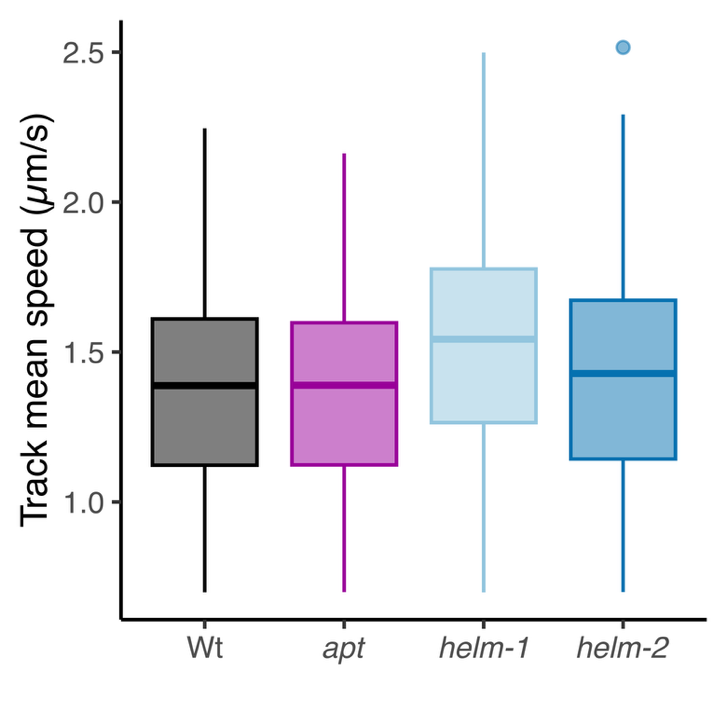

**Figure S2: Swimming speed of of Wt and helm gametes.** Box plot of the average speed along the tracked trajectories (n>100 trajectories).

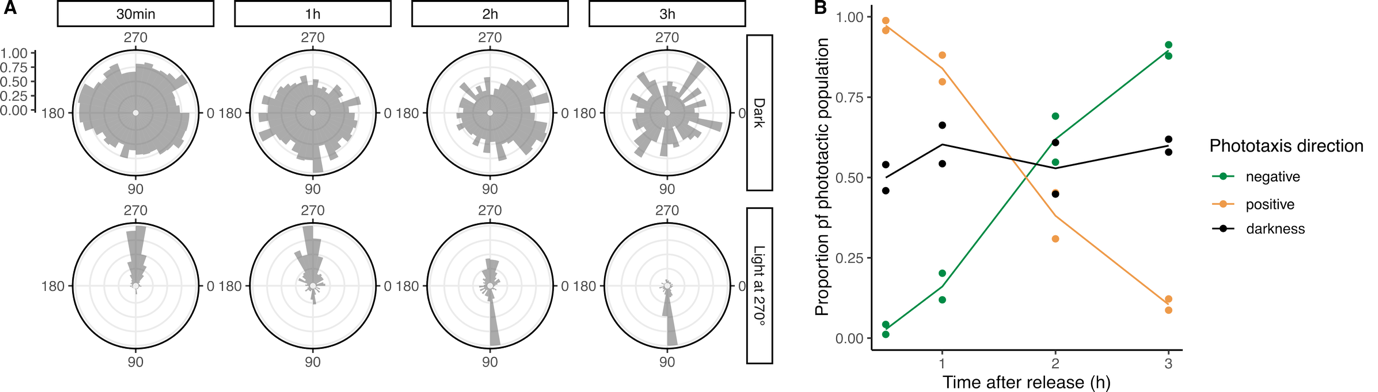

**Figure S3: Change in the phototactic sign of Ectocarpus wild-type gametes with time after their release*.*** (A), Polar histograms of swimming direction under different light regimes (counts binned by 10° bin width). (B), Proportion of negatively and positively phototactic gametes with time after the gametes release. Black lines and dots represent the proportion of gametes swimming in the 90° direction in the dark. Two technical replicates are represented.

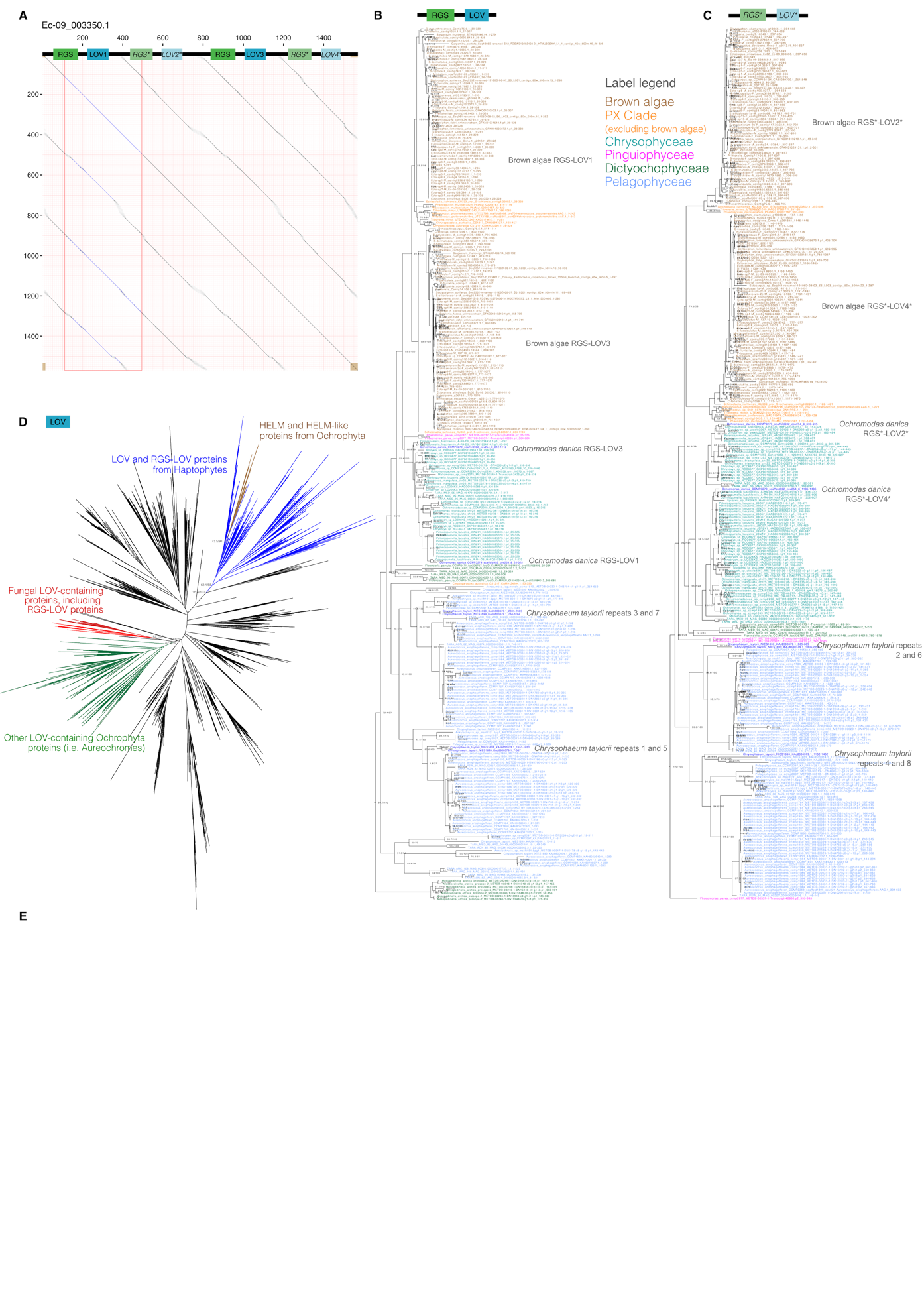

**Figure S4. Evolutionary history of HELM and HELM-like proteins.** *(A), Dot plot analysis of the Ectocarpus HELM protein, revealing the repeat structure of the protein, generated with HHrepID^35,36^. B and C, Phylogenetic tree of the functional (B) and non-functional (C) RGS-LOV repeats. (D), phylogenetic tree of the LOV domain, showing the deep branching of the HELM and HELM-like proteins within proteins for Haptophytes.*

**Supplementary Video 1**: Photoaccumulation of Wt gametes

**Supplementary Video 2**: Tracking of swimming Wt gametes in the dark. Each track retained for analysis is randomly colored and fades in 0.5s.

**Supplementary Video 3**: Tracking of swimming Wt gametes under unidirectional light from the top

**Supplementary Video 4**: Tracking of swimming *helm*-1 gametes under unidirectional light from the top
